# Morphogenesis of the islets of Langerhans is guided by extra-endocrine Slit2/3 signals

**DOI:** 10.1101/2020.06.15.153353

**Authors:** Jennifer M. Gilbert, Melissa T. Adams, Nadav Sharon, Hariharan Jayaraaman, Barak Blum

**Author notes:** Corresponding author: Barak Blum.

## Abstract

The spatial architecture of the islets of Langerhans is vitally important for their correct function, and alterations in islet morphogenesis often result in diabetes mellitus. We have previously reported that Roundabout (Robo) receptors are required for proper islet morphogenesis. As part of the Slit-Robo signaling pathway, Robo receptors work in conjunction with Slit ligands to mediate axon guidance, cell migration, and cell positioning in development. However, the role of Slit ligands in islet morphogenesis has not yet been determined. Here we report that Slit ligands are expressed in overlapping and distinct patterns in both endocrine and non-endocrine tissues in late pancreas development. We show that function of either Slit2 or Slit3, which are predominantly expressed in the pancreatic mesenchyme, is required and sufficient for islet morphogenesis, while Slit1, which is predominantly expressed in the β cells, is dispensable for islet morphogenesis. We further show that Slit functions as a repellent signal to β cells. These data suggest that clustering of endocrine cells during islet morphogenesis is guided, at least in part, by repelling Slit2/3 signals from the pancreatic mesenchyme.

## Introduction

Blood glucose homeostasis is regulated in the pancreas by clusters of endocrine cells called the islets of Langerhans. Islets consist of five different endocrine cell types (α, β, δ, PP, ε), which secrete glucagon, insulin, somatostatin, pancreatic polypeptide, and ghrelin, respectively. Murine islets exhibit a distinct cytoarchitecture consisting of a core of β-cells, surrounded by a mantle of α-, δ-, PP- and ε-cells. The β-cell core makes up roughly 80% of the islet mass, while the four other cell types make up the remaining 20% (Kim et al., 2009; Steiner et al., 2010). This cytoarchitecture is thought to be important for proper islet function, and loss of this architectural makeup is described in obesity and diabetes in both mice and humans (Baetens et al., 1978; Cabrera et al., 2006; Kilimnik et al., 2011; Roscioni et al., 2016). While the architectural features of islets have been well-documented, the mechanisms controlling the formation this architecture are still largely unknown.

The Slit-Robo signaling pathway has roles in a number of developmental processes, primarily axon guidance, cell movement, and cell adhesion (Blockus and Chédotal, 2016; Chédotal, 2007; Wu et al., 2017; Ypsilanti and Chedotal, 2014; Ypsilanti et al., 2010). Slit ligand binding to Robo receptors can induce cell migration using repulsive or attractive cues in a context-dependent manner. In the developing mouse, Slit-Robo signaling provides a repulsive corridor to prevent migrating axons from straying from their path during innervation (Brose et al., 1999; Dickson and Gilestro, 2006). Slit-Robo binding inactivates Rho GTPases, inhibiting actin polymerization and driving the cell away from the direction of the Slit signal (Wu et al., 2017; Ypsilanti et al., 2010). Conversely, Slit uses attractive cues to promote vascular development and angiogenesis. In this context, Slit-Robo interactions activate Rho GTPases, inducing actin polymerization in the direction of the Slit signal (Rama et al., 2015; Wu et al., 2017; Ypsilanti et al., 2010; Zhang et al., 2009). While Slit and Robo are a canonical signaling pair, both components have alternative binding partners; Slit ligands are able to bind semaphorins, ephrins, plexin, and neuronatin to regulate cell migration and metabolic function in specific tissues (Brose et al., 1999; Delloye-Bourgeois et al., 2015; Svensson et al., 2016; Wright et al., 2012). Robo receptors are able to bind the fibronectin leucine rich transmembrane protein 3 (FLIRT3), and are capable of forming homodimers to induce axonal growth (Hivert, 2002; Leyva-Díaz et al., 2014; Tong et al., 2019).

We have recently described a role for Robo receptors in pancreatic islet architecture (Adams et al., 2018). Specifically, we showed that genetic deletion of *Robo1* and *Robo2* in β-cells (*Robo* β*KO*) results in loss of stereotypic murine islet architecture, without affecting β-cell differentiation or maturation. These Robo-depleted islets have a marked invasion of α- and δ-cells into the β-cell core. Given the conserved role of Slits as the canonical Robo ligands and our recent findings that Robo receptors regulate endocrine cell type sorting in the islet, we set to investigate the role of Slit ligands in islet morphogenesis.

## Results

### Slits ligands are expressed in different compartments in the developing mouse pancreas

To test the hypothesis that Slits are involved in Robo-mediated control of islet architecture during development, we first examined whether any of the Slit ligands are expressed in the pancreas at the time of islet morphogenesis. We queried a gene expression database, generated by Krentz and colleagues (Krentz et al., 2018), which contains single-cell RNA-Seq data from embryonic mouse pancreata. We found that *Slit1* expression is present in a subset of endocrine progenitor cells at embryonic day (E)15.5, and becomes enriched in β-cells by E18.5. *Slit2* and *Slit3* expression is distributed between pancreatic mesenchyme, acinar, and ductal cell types with negligible expression in the endocrine compartment at both time points (Figure 1).

**Figure 1:**
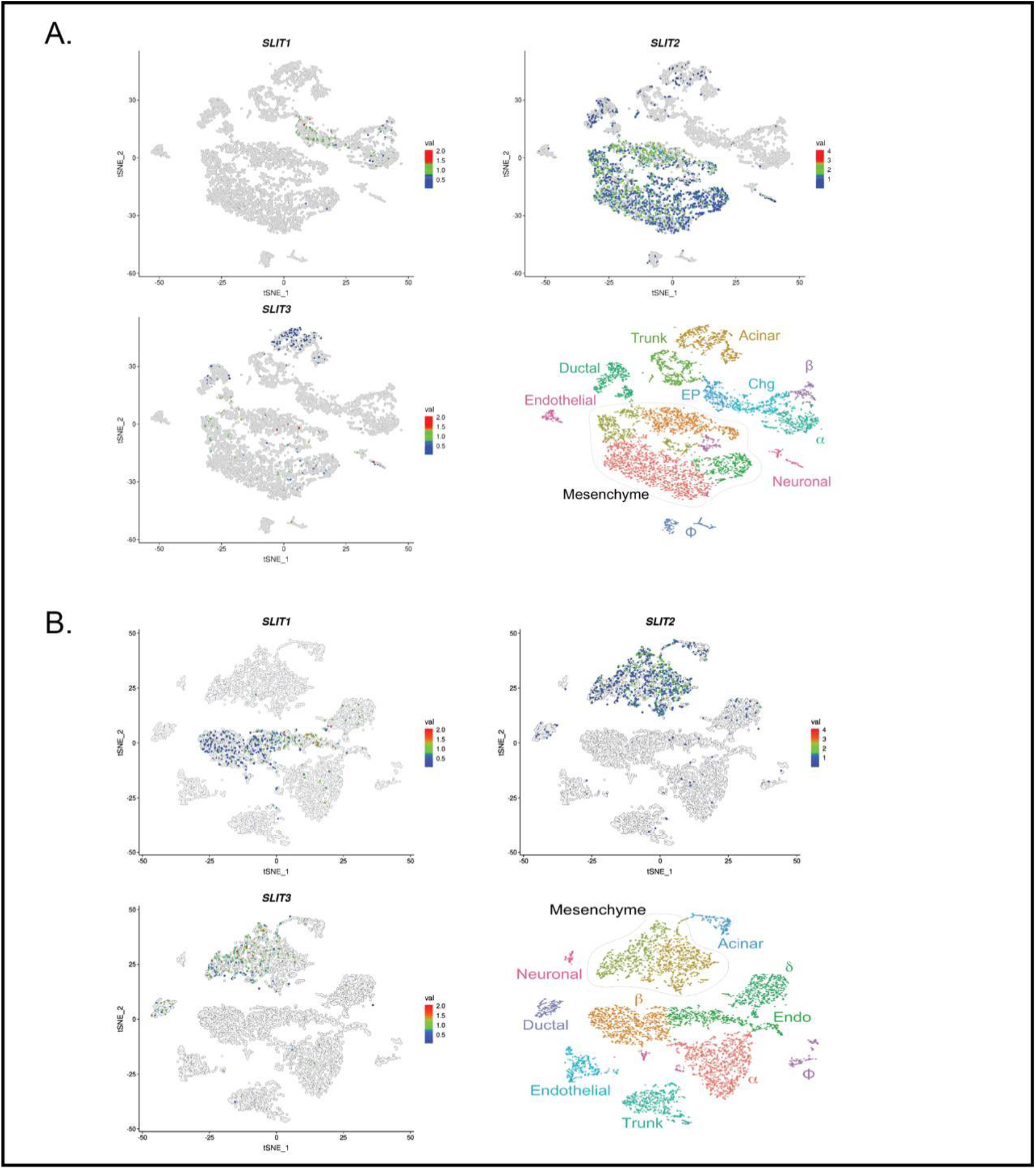
Slit transcripts are expressed in different compartments in the developing murine pancreas. Single-cell RNA-Seq data (scRNA-Seq) adapted from Krentz and colleagues (Krentz et al. 2018). tSNE plots depicting Slit1, Slit2, and Slit3 expression in pancreatic cells. Time points analyzed are E15.5 (A) and E18.5 (B). Slit1 is restricted to the endocrine compartment, while Slit2 and Slit3 localize with the mesenchyme/acinar compartment.

To confirm the expression of Slits in the pancreas *in vivo*, we analyzed pancreata from *Slit1*^*GFP*^, *Slit2*^*GFP*^, and *Slit3*^*LacZ*^ mice, which have knock-in reporters at their respective endogenous Slit loci (Plump et al., 2002; Yuan et al., 2003). We identified strong GFP expression in *Slit1*^*GFP*/+^ mice both in E18.5 and adult islets. This staining pattern overlapped with insulin, indicating that *Slit1* is expressed in β-cells at both stages (Figure 2A). We did not detect *Slit2*^*GFP*^ (Figure 2A) or *Slit3*^*LacZ*^ (Figure 2B) in either the embryonic or the adult islets. However, *Slit3*^*LacZ*^ expression was detected in pancreatic tissues outside of the islet (Figure 2B). *Slit2*^*GFP*^ expression was seen in other tissues, indicating that the lack of *Slit2*^*GFP*^ signal in the developing pancreas is not caused by a problem with the reporter (Supplementary Figure 1). A previous report by Escot and colleagues identified *Slit3* expression in the developing pancreatic mesenchyme (Escot et al., 2018). While we were not able to detect *Slit2*^*GFP*^ expression, data from single-cell RNA Sequencing (scRNAseq) indicates that it is also expressed in pancreatic mesenchyme during development (Krentz et al., 2018). We concluded that *Slit1* is the predominant Slit expressed inside the islets, and that *Slit3* and perhaps *Slit2* are expressed outside of the islet during pancreatic development.

**Figure 2:**
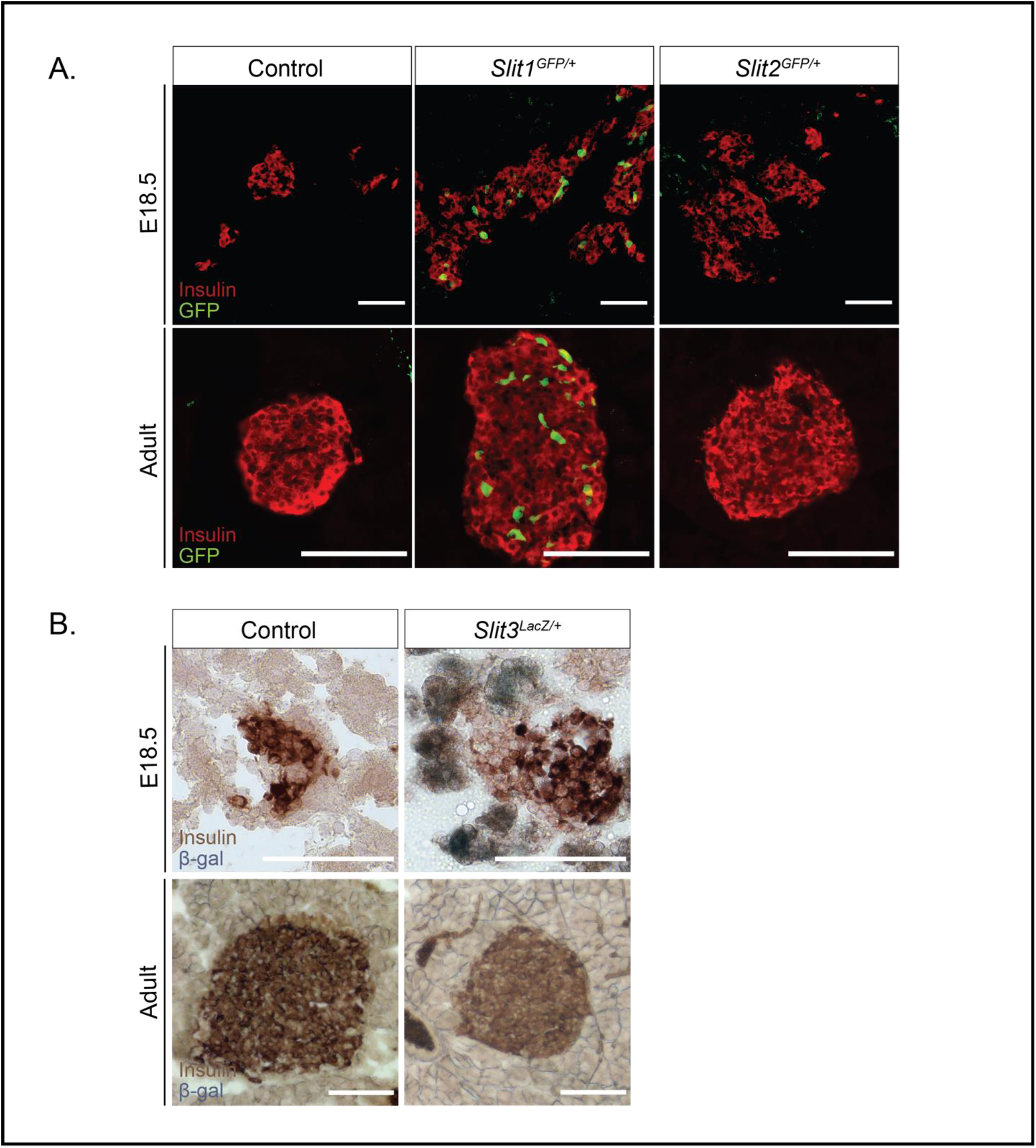
Slit1, but not Slit2 or Slit3, is expressed in the mouse islet from embryonic stages to adulthood. (A) Immunofluorescence staining of β-cells (Insulin, red) and Slit1, Slit2 (GFP, green) in E18.5 and adult heterozygous knock-in mice. (B) β-gal staining of Slit3 (LacZ, blue) in E18.5 and adult heterozygous knock-in mice. β-gal staining (Slit3 expression) is apparent in non-endocrine tissue surrounding the islet in the embryo. Scale bar = 100 microns.

### Loss of a single Slit ligand does not compromise islet architecture

Slit and Robo are conserved binding partners, and loss of Robo in the islets of *Robo* β*KO* mice results in severely altered islet architecture (Adams et al., 2018). We hypothesized that if Slits mediate Robo-regulated islet architecture, then eliminating Slit expression would phenocopy the islet organization defects in *Robo* β*KO* islets. Whole-body *Slit1*-null (*Slit1*^*GFP/GFP*^) and *Slit3*-null (*Slit3*^*LacZ/LacZ*^) mice are viable to adulthood. We performed positional cell counting on the islets of these mice as previously described (Adams et al., 2018) to determine whether these mutants exhibited islet organizational defects. In contrast to the phenotype seen in *Robo* β*KO* islets, individual *Slit1* or *Slit3* mutant islets display completely normal architecture (Figure 3A-C). α-, β-, and δ-cells remain restricted to their respective niches; the β-cells reside in the core, while the α- and δ-cells remain in the islet mantle. We also found no significant difference between control islets and *Slit1* or *Slit3* mutant islets in islet size (Figure 3D) or circularity (Figure 3E). Whole-body *Slit2*-null (*Slit2*^*GFP/GFP*^) animals die shortly after birth. We thus examined pancreata of E18.5 *Slit2*^*GFP/GFP*^ embryos. Evidence of altered islet architecture in *Robo* β*KO* mutants can be seen at E18.5; however, we did not observe overt defects in the architecture of *Slit2*^*GFP/GFP*^ islets at this time point (Figure 3F). Taken together, these results indicate that individual Slits are either not required for, or compensate for each other in, Robo-mediated control of islet architecture.

**Figure 3:**
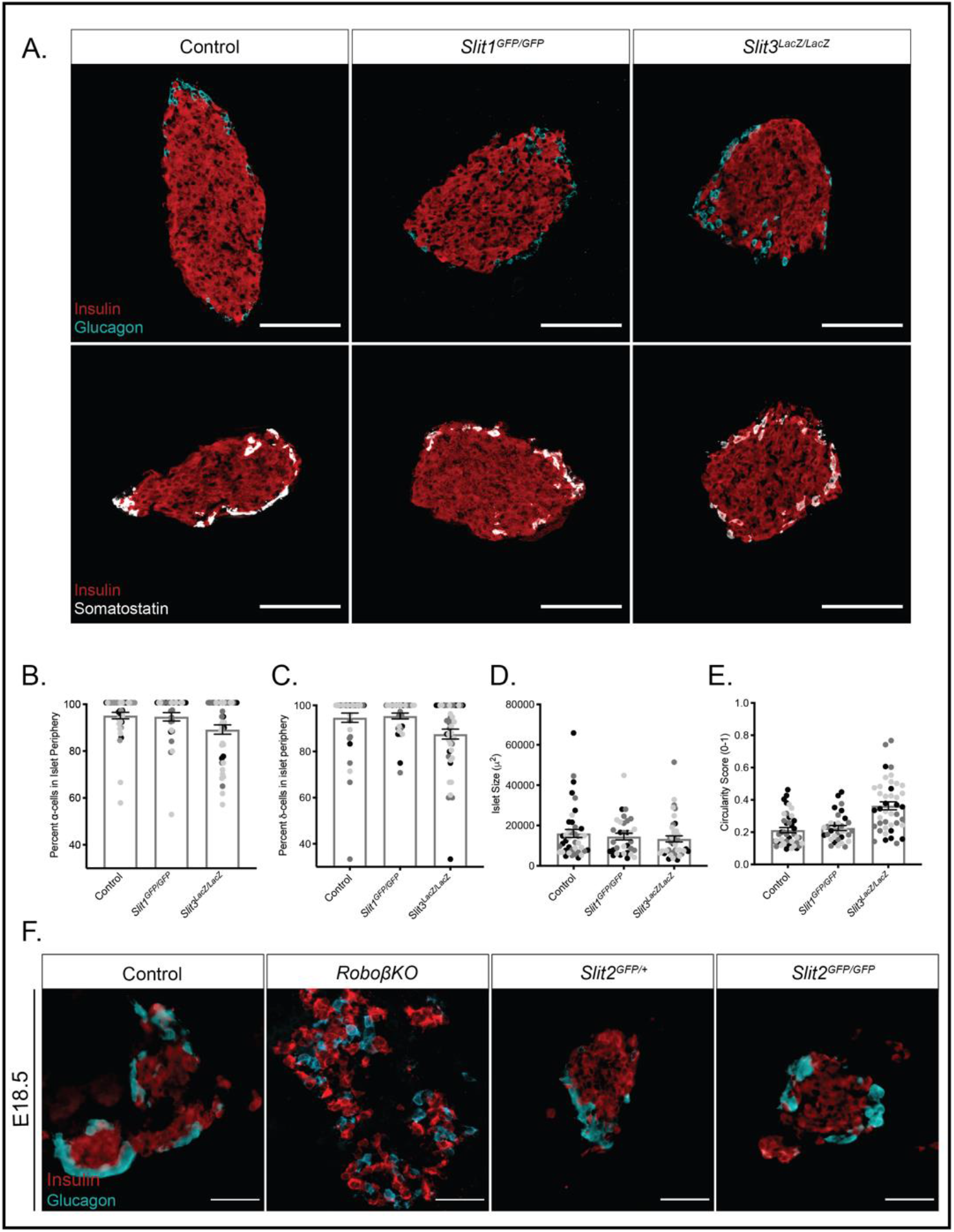
Loss of a single Slit ligand does not compromise islet architecture. (A) Immunofluorescence staining of β-cells (Insulin, red), α-cells (Glucagon, cyan) and δ-cells (Somatostatin, white) in adult (~8 week old) homozygous knockout mice. Scale bar = 100 microns. (B) Percentage of α-cells found in the islet periphery out of total α-cells. (C) Percentage of δ-cells found in the islet periphery out of total δ-cells. (D) Average islet size (E) Average islet circularity (as noted by a circularity score of 0-1, where 1 is a perfect circle). (F) Immunofluorescence staining of β-cells (Insulin, red) and α-cells (Glucagon, cyan) in E18.5 control, Ins2-Cre;Robo1^−/^;Robo2^flx/flx^, and Slit2 mice. Scale bar = 50 microns. Data presented as mean ±SEM.

### Slit2 and Slit3 compensate for each other and are required for islet morphogenesis

Slit ligands are highly similar in amino acid sequence, particularly in their Robo-binding domains (Figure 4A). Thus, it is possible that the different Slit ligands compensate for each other during islet morphogenesis. We tested the extent to which multiple Slit ligands are required for islet architecture by analyzing islet formation in combinatorial Slit mutants. *Slit1*^*GFP/GFP*^;*Slit3*^*LacZ/LacZ*^ double knockouts live to adulthood and appear normal, with no detectable alterations in islet architecture, size, or circularity (Figure 4B-F). To circumvent the neonatal lethality of *Slit2*^*GFP/GFP*^ mice, we analyzed the pancreata of *Slit1/2* knockouts (*Slit1*^*GFP/GFP*^;*Slit2*^*GFP/GFP*^), *Slit2/3* knockouts (*Slit2*^*GFP/GFP*^;*Slit3*^*LacZ/LacZ*^), and *Slit1/2/3* knockouts (*Slit1^GFP/GFP^;Slit2^GFP/GFP^;Slit3^LacZ/LacZ^*) at E18.5 or P0. *Slit1/2* knockout islets show no indications of altered architecture, but *Slit2/3* and *Slit1/2/3* knockouts have disorganized islets (Figure 4G). To quantify this phenotype, we scored islets as either intact (insulin-positive cells surrounded by glucagon-positive cells), intermediate (clusters of insulin-positive cells disrupted by glucagon-positive or non-endocrine cells), or disrupted (single cells or clusters of endocrine cells that are not forming islet structures) (Figure 4H). Double-blinded scoring of islets from the above genotypes revealed that wild type and *Slit1/2* knockouts have few disrupted islets and similar percentages of intact and intermediate islets (*WT* intact: 49%, intermediate: 40%, disrupted: 11%. *Slit1/2 KO* intact: 43%, intermediate: 46%, disrupted: 11%). On the other hand, *Slit2/3* and *Slit1/2/3* knockouts had very few intact islets and increased numbers of intermediate and disrupted islets (*Slit2/3 KO* intact: 8%, intermediate: 53%, disrupted: 39%. *Slit1/2/3* KO intact: 8%, intermediate: 60%, disrupted: 32%). Taken together, the data suggest that *Slit1* (expressed in the islet itself) is dispensable, while *Slit2* and *Slit3* (expressed outside of the islet) compensate for each other and are required for proper islet formation.

**Figure 4:**
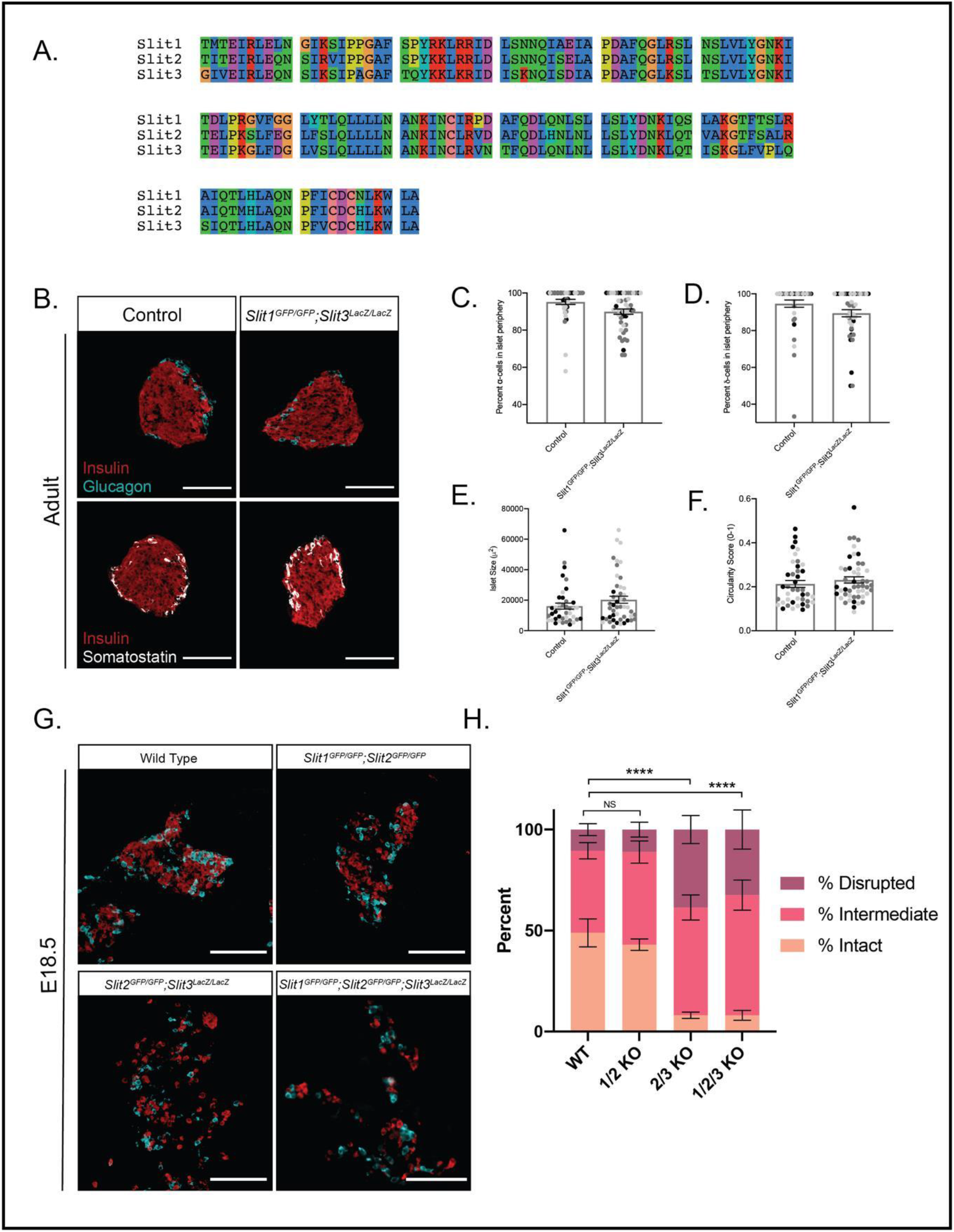
Slit2 and Slit3 compensate for one another in islet morphogenesis. (A) Amino acid alignment of the Robo-binding domain LRR2 (Slit1: 310aa-451aa, Slit2: 301aa-442aa, Slit3: 308aa-449aa) of all three murine Slits. In this region, the pairwise alignment scores are: Slit1/Slit2: 77%, Slit1/Slit3: 69%, Slit2/Slit3: 70%. (B) Immunofluorescence staining of β-cells (Insulin, red), α-cells (Glucagon, cyan) and δ-cells (Somatostatin, white) in adult (~8 week old) homozygous Slit1^GFP/GFP^;Slit3^LacZ/LacZ^ knockout mice. Scale bar = 100 microns. (C) Percentage of α-cells found in the islet periphery out of total α-cells. (D) Percentage of δ-cells found in the islet periphery out of total δ-cells. (E) Average islet size (F) Average islet circularity (as noted by a circularity score of 0-1, where 1 is a perfect circle). Data presented as mean ±SEM. (G) Immunofluorescence staining of β-cells (Insulin, red) and α-cells (Glucagon, cyan) in control (wild-type), double, and triple knockout mice at E18.5/P0. Scale bar = 100 microns. (H) Percentage of islets from each genotype that were scored as intact, intermediate, or disrupted. Data presented as mean ±SEM. p <0.0001; Chi-square test.

### Slits act as repellent factors to influence β cell migration

Because *Slit2/3* and *Slit1/2/3* mutant islets are disrupted and do not cluster tightly, we wondered whether this indicates failure of β-cells to migrate properly during islet morphogenesis. To test this hypothesis, we performed Transwell cell migration assays using INS-1 cells. INS-1 cells seeded in the top chamber of a cell culture insert above INS-1 conditioned media showed strong migratory activity, while INS-1 cells seeded above fresh, untreated INS-1 culture media did not (Figure 5A-C). INS-1 cells seeded above conditioned media supplemented with 2.5μg recombinant SLIT1, SLIT2, and SLIT3 displayed a significantly reduced ability to migrate (Figure 5B,C), suggesting that Slits influence β-cell migration through cell-cell repulsion mechanisms during islet morphogenesis.

**Figure 5:**
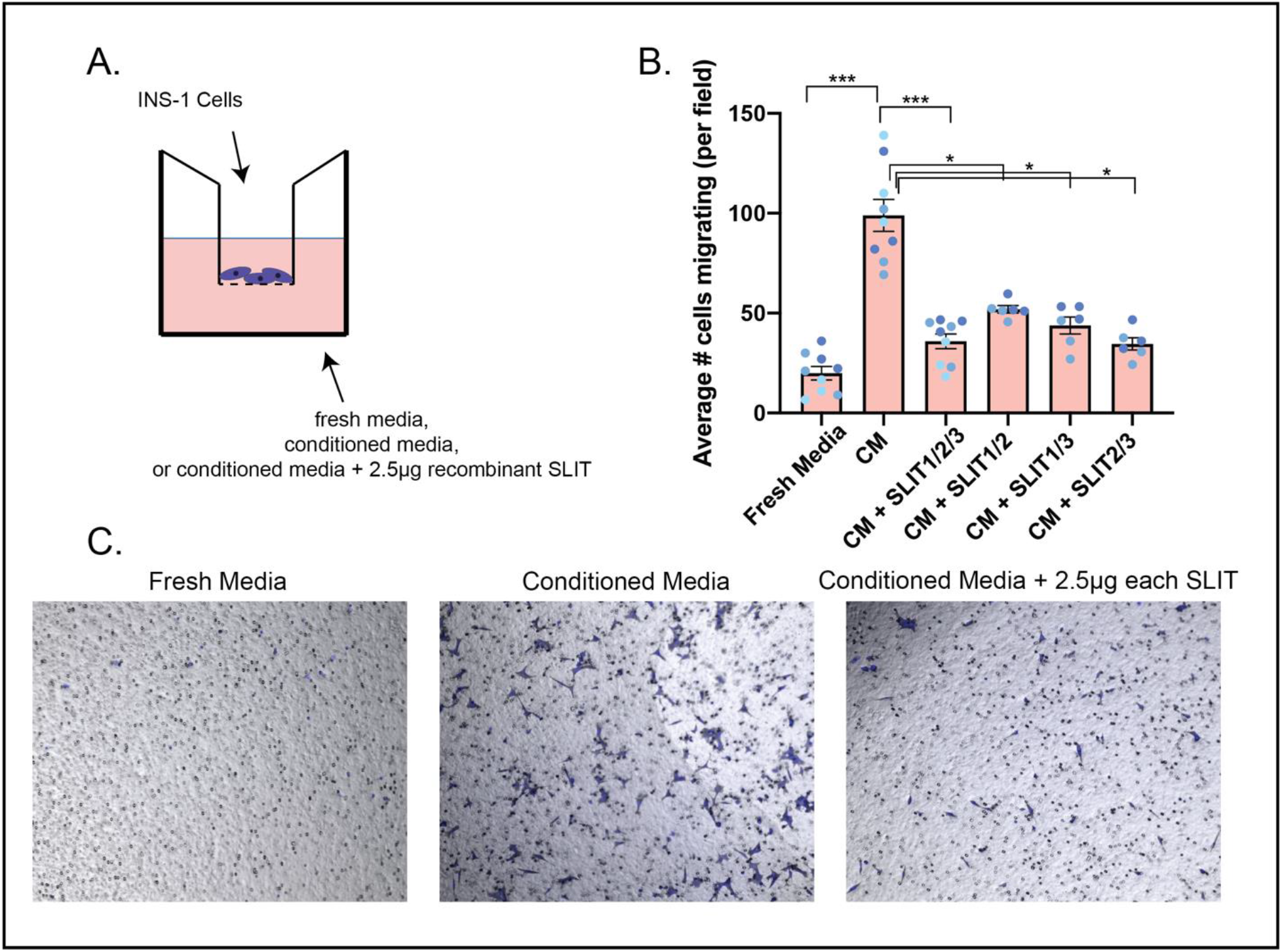
Slits act as repellent factors to influence β-cell migration. (A) Schematic diagram of Transwell cell migration assay. INS-1 cells were seeded in cell culture inserts over INS-1 conditioned media, fresh culture media, or INS-1 conditioned media supplemented with 2.5μg of each recombinant SLIT protein. (B) Results of cell migration assay. The average number of cells migrating per field of view is plotted. Data presented as mean ±SEM. ***p<0.0005 *p<0.05 (C) Representative images of a single field of view of a cell migration insert used in the experiments shown in (B).

## Discussion

In this study, we demonstrate that Slit ligands are required for pancreatic islet architecture. Simultaneous loss of all three Slits results in a disrupted, “islet explosion-like” phenotype, which is also observed in *Slit2*^*GFP/GFP*^;*Slit3*^*LacZ/LacZ*^ knockouts. These findings lead us to conclude that Slit1 is dispensable for, and that *Slit2* and *Slit3* are required for and have redundant roles in, islet morphogenesis.

The exact mechanism of Slits in islet morphogenesis is unknown; however, the expression pattern of Robo in the pancreas provides some clues. Slit and Robo are ligand-receptor binding partners in the Slit-Robo signaling pathway. During mammalian development, Slit and Robo occupy adjacent tissues, specifying complimentary expression patterns in the developing organism (Yuan et al., 1999). While all three Slits have expression patterns unique to their specific domain, they also have overlapping regions of expression, suggesting some genetic redundancy. Interestingly, *Slit2* and *Slit3* share more expression domains with each other than either of them do with *Slit1* (Yuan et al., 1999). We have observed a similar framework in the mouse pancreas: *Robo* is primarily expressed in endocrine cells (Adams et al., 2018), while *Slit2* and *Slit3* have overlapping expression patterns in pancreatic mesenchyme. These complimentary and overlapping regions of expression are hallmarks of ligand-receptor binding partners, and suggest that mesenchymal Slit2 and Slit3 interact with endocrine Robo to coordinate islet morphogenesis. We propose that Slit2/3 signals from the mesenchyme are picked up by Robo receptors on the surface of developing islet endocrine cells. These endocrine cells are then repelled away from the direction of the Slit signal, allowing for islet clustering to occur. Loss of this signal results in a failure of islet morphogenesis, thus the “islet explosion” phenotype described above. It is likely that Slits are not the only signal required for morphogenesis, as some of the islets in *Slit2*^*GFP/GFP*^;*Slit3*^*LacZ/LacZ*^ and triple knockout animals showed evidence of appropriate clustering. Future work will determine whether other ligands or even Robo-Robo interactions are involved in islet morphogenesis.

It is commonly held that islet morphogenesis is outlined by delamination of endocrine progenitors from the pancreatic duct, followed by their migration as individual cells though the mesenchyme and aggregation into islets (Pan and Wright, 2011), implying that β-cells respond to attractive cues from the islets. Indeed, we have observed strong Transwell migration of β-cells towards their own conditioned medium, demonstrating that β-cells are attracted towards β-cells. However, we further provide evidence to suggest mesenchymal Slits repel β-cells during islet morphogenesis. These results are in support of the recent observation that endocrine progenitors remain physically connected throughout islet morphogenesis (Sharon et al., 2019), and suggest that after β-cell delamination, repulsion of the β-cells by mesenchymal Slits pushes them into the center of the islet, thus maintaining the core-mantle architecture. Taken together, we propose that both attractive and repulsive signals operate together in forming the canonical murine islet architecture.

The role of *Slit1* in islet morphogenesis remains elusive. The islet architecture phenotypes seen in *Slit2*^*GFP/GFP*^;*Slit3*^*LacZ/LacZ*^ animals are not significantly different in triple knockouts, suggesting that *Slit1* does not have any influence on islet architecture. In addition, *Slit1* expression does not overlap with *Slit2* or *Slit3*, thus it is unlikely to be redundant. Morover, *Slit1* is strongly expressed in a subset of β-cells both during development and in the adult. Slits have been previously demonstrated to provide a protective effect on β-cells, as well as potentiate insulin secretion (Yang et al., 2013). It is thus plausible that *Slit1* is tasked with the protective and secretory roles in the islet, while the transient expression of *Slit2* and *Slit3* in the mesenchyme during development is responsible for islet morphogenesis. How the expression of *Slit1* in β-cells does not interfere with the function of Slit2 and Slit3 from the mesenchyme during islet morphogenesis is intriguing, and remains to be elucidated.

## Methods

### Animals

All animal experiments were conducted in accordance with the University of Wisconsin-Madison IACUC guidelines under approved protocol #M005221. *Robo1*^−^;*Robo2*^*flx*^ (Branchfield et al., 2016), *Ins2-Cre* (Postic et al., 1999), *H2B-mCherry* (Blum et al., 2014), *Slit1*^*GFP*^, *Slit2*^*GFP*^ (Plump et al., 2002), *Slit3*^*LacZ*^ (Yuan et al., 2003) alleles have been previously described.

### Expression Analysis

tSNE plots of sc-RNA Seq obtained from the Lynn Lab’s Single Cell Gene Expression Atlas (https://lynnlab.shinyapps.io/embryonic_pancreas/) (Krentz et al., 2018).

### Immunostaining

Pancreata were dissected from adult (~8week old), embryonic (E18.5), or newborn (P0) mice, fixed in 4% paraformaldehyde for 1 hour at room temperature (20-30 minutes for E18.5 and P0), preserved in 30% sucrose, embedded in OCT (Leica), then sectioned onto slides. Slides were stained according to the following protocol: 1 hour block in 10% Normal Donkey Serum in PBST, 1 hour primary antibody incubation, 3×10 min PBST washes, 1 hour secondary antibody incubation in dark, 3×10 min PBST washes, mount slides in Fluoromount-G (Thermo Fisher). The following primary antibodies were used: Guinea Pig anti-Insulin 1:800 (Dako), Guinea Pig anti-Insulin pre-diluted 1:6 (Dako 1R002), Chicken anti-GFP 1:1000 (Abcam ab13970), Rabbit anti-Glucagon 1:200 (Cell Signaling 2760), Goat anti-Somatostatin 1:50 (Santa Cruz), Rabbit anti-Somatostatin 1:800 (Phoenix G-060-03), DAPI 1:10,000 (Sigma 9542). The following secondary antibodies were used at 1:500: Alexa 647 anti-Guinea Pig, Alexa 594 anti-Rabbit, Alexa 594 anti-Goat, Alexa 488 anti-Rabbit, Alexa 488 anti-Chicken.

For eye analysis, tissues were dissected and fixed in 4% paraformaldehyde for 2 hours at 4°C. Tissues were preserved in a series of sucrose solutions (10%, 20% sucrose) for 1.5 hours each. Tissues were further preserved in 30% sucrose overnight, embedded in OCT, then sectioned and stained as above. For β-galactosidase staining, tissues were fixed in 4% paraformaldehyde for 1 hour at room temperature (or 20-30 minutes for E18.5 tissue). Fixed tissues were stained with X-gal solution (Roche 11828673001) for 22 hours at 37°C, then preserved, embedded, and sectioned as above. Insulin staining on these tissues was done using the Vectastain ABC HRP kit (Vector Labs PK-4007), NovaRED kit (Vector Labs SK-4800), and mounted with VectaMount (Vector Labs H-5000). Slides for expression analysis imaged using a Zeiss Axio Observer Z1.

### Cell Counting, Shape, Size Analysis

Slides used for cell counting or shape and size analysis were imaged on a Nikon A1RS confocal microscope. Confocal z-stacks were converted to maximum intensity projected images. The number of α- and δ-cells were counted using the ImageJ Cell Counter tool. α- or δ-cells were considered in the islet periphery if they were within the first two cell layers of the islet. For shape and size analysis, islets were outlined and a threshold was applied in ImageJ. The Analyze Particles tool then gave readout of islet size in μm^2^ and a circularity score (between 0-1, where 1 indicates a perfect circle). A minimum of 10 islets were analyzed across at least three different tissue sections per mouse. Analysis performed on *n=3* mice for each genotype. α- and δ-cell percentages, islet size, and islet circularity values were averaged for each mouse and plotted in Prism.

### Amino Acid Alignment

Amino acid sequence and domain information were obtained from Yuan et al., 1999. Pairwise alignment scores of amino acid sequences were provided by ClustalW (https://www.genome.jp/tools-bin/clustalw).

### Islet Scoring

Islet scoring was performed on images of tissue sections stained for insulin, glucagon, and DAPI. Z-stack images were converted to maximum intensity projected images, and randomly assigned a number identifier. Four independent trials (by four different researchers) of double-blinded scoring was performed on 197 images, comprising at least 10 images spanning four different tissue sections per mouse, and at least 3 mice per genotype.

### Transwell Cell Migration Assay

INS-1 cells (AddexBio) were maintained in culture media containing RPMI-1640 (ThermoFisher), 10% FBS, 1% penicillin/streptomycin, and supplemented with 0.05mM β-mercaptoethanol. Cells were seeded at a density of 250,000 cells/mL in Transwell cell culture inserts with 8μM pores (Sigma). Inserts were placed into wells containing either 700μL culture media, 700μL INS-1 conditioned media, or 700μL INS-1 conditioned media supplemented with 2.5μg each recombinant SLIT1, SLIT2, and SLIT3 (R&D Systems) and cultured at 37°C for 48 hours. Inserts were then fixed in 4% paraformaldehyde for 20 minutes, unmigrated cells were wiped off the top of the insert, and then inserts were incubated in 0.08% crystal violet and a 1:1,000 concentration of DAPI to visualize the cells. Nine non-overlapping field of view images were taken for each insert. Three images per insert were chosen at random for quantification. Results are reported as the average number of cells that migrated per field of view.

### Statistical Analysis

All data reported as mean ±SEM unless otherwise indicated. P-values calculated using Student’s T-test in Prism GraphPad 7 unless otherwise indicated. Any p-value <0.05 was considered significant and marked with an asterisk.

## Author Contributions

Conceptualization, B.B. and J.M.G; Methodology, B.B. and J.M.G; Investigation, J.M.G, M.T.A., N.S, and H.J.; Formal Analysis, J.M.G, M.T.A., N.S, and H.J.; Writing Original Draft, B.B and J.M.G.; Writing, Review and Editing, all authors; Funding Acquisition, B.B.; Supervision, B.B.

## Acknowledgements

We thank members of the Blum lab, especially Bayley Waters and Dex Nimkulrat for valuable discussion and comments on the manuscript. We thank Le Ma, David Ornitz, Alain Chedotal, and Marc Tessier-Lavigne for mice. We are grateful to Francis Lynn and Nicole Krentz for allowing us to use their scRNA-seq data, and to Cody Frederickson for help generating figures. We are also grateful to Lance Rodenkirch and the UW-Madison Optical Imaging Core for help with imaging. This work was funded in part by the following grants. R01DK121706 from the NIDDK, the DRC at Washington University Pilot Grant P30DK020579, and Pilot Award UL1TR000427 from the UW-Madison Institute for Clinical and Translational Research (ICTR). JMG and MTA were funded by 5T32GM007133-44, a graduate training award from the UW-Madison Stem Cell & Regenerative Medicine Center, and an Advanced Opportunity Fellowship through SciMed Graduate Research Scholars at UW-Madison.

---

**Supplemental Figure 1:**
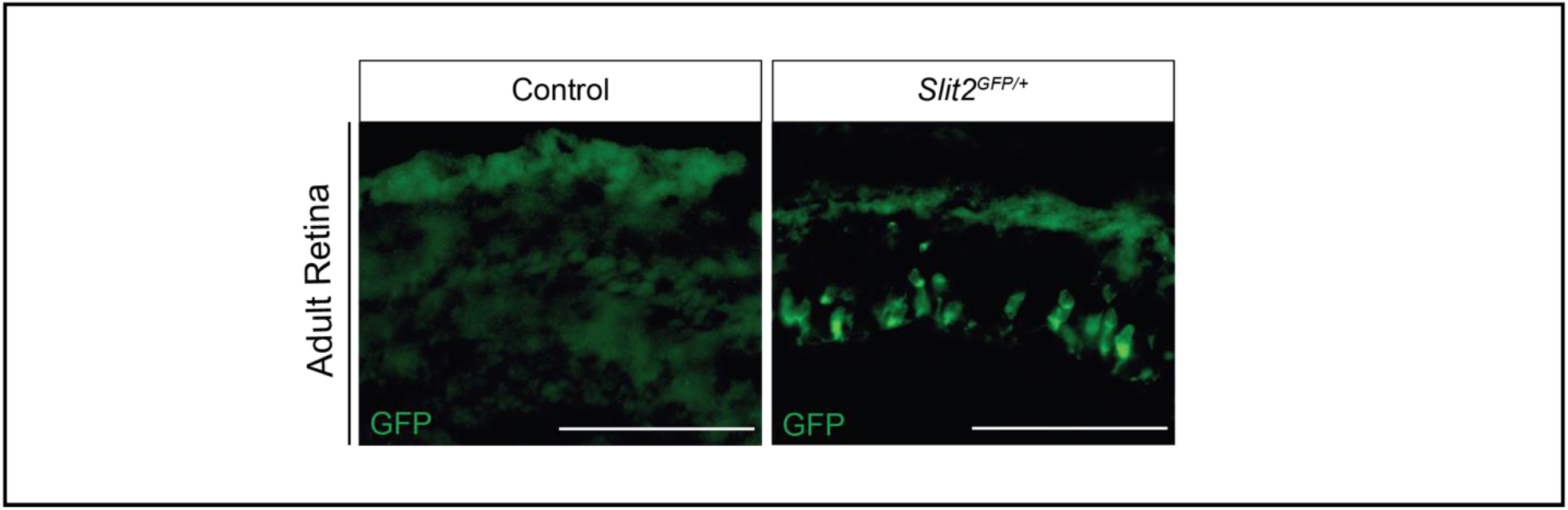
The Slit2^GFP^ reporter is functional. Immunofluorescence staining of Slit2 (GFP, green) in retinal sections from control (wild type) or Slit2^GFP/+^ heterozygous animals. Scale bar = 100 microns.

